# Deep learning-based real-time detection of novel pathogens during sequencing

**DOI:** 10.1101/2021.01.26.428301

**Authors:** Jakub M. Bartoszewicz, Ulrich Genske, Bernhard Y. Renard

## Abstract

Novel pathogens evolve quickly and may emerge rapidly, causing dangerous outbreaks or even global pandemics. Next-generation sequencing is the state-of-the-art in open-view pathogen detection, and one of the few methods available at the earliest stages of an epidemic, even when the biological threat is unknown. Analyzing the samples as the sequencer is running can greatly reduce the turnaround time, but existing tools rely on close matches to lists of known pathogens and perform poorly on novel species. Machine learning approaches can predict if single reads originate from more distant, unknown pathogens, but require relatively long input sequences and processed data from a finished sequencing run. Incomplete sequences contain less information, leading to a trade-off between sequencing time and detection accuracy. Using a workflow for real-time pathogenic potential prediction, we investigate which subsequences already allow accurate inference. We train deep neural networks to classify Illumina and Nanopore reads and integrate the models with HiLive2, a real-time Illumina mapper. This approach outperforms alternatives based on machine learning and sequence alignment on simulated and real data, including SARS-CoV-2 sequencing runs. After just 50 Illumina cycles, we observe an 80-fold sensitivity increase compared to real-time mapping. The first 250bp of Nanopore reads, corresponding to 0.5s of sequencing time, are enough to yield predictions more accurate than mapping the finished long reads. The approach could also be used for screening synthetic sequences against biosecurity threats.

## Introduction

### Background

The SARS-CoV-2 coronavirus emerged in late 2019 causing an outbreak of COVID-19, a severe respiratory disease, which quickly developed into a global pandemic of 2020. This virus of probable zoonotic origin [1] is a terrifying example of how easily new agents can spread. What is more, many more novel pathogens are expected to emerge. They evolve extremely quickly due to high mutation rates or horizontal gene transfer, while human exposure to the vast majority of unexplored microbial biodiversity is rapidly growing [2, 3]. New biosafety threats may come as novel bacterial agents like the Shigatoxigenic *Escherichia coli* strain that caused a deadly epidemic in 2011 [4]. Outbreaks of previously unknown viruses can be even more severe. This is not limited to coronaviruses like SARS-CoV-1, MERS and SARS-CoV-2. Novel strains of the Influenza A virus caused four pandemics in less than a hundred years (from the “Spanish flu” of 1918 until the “swine flu” of 2009), killing millions of people. Even known pathogens may be difficult to control, as proven by the outbreaks of Zika and Ebola in the 2010s [5]. Importantly, many viruses can switch between more than one host or evolve silently in an animal reservoir before infecting humans. This happened before to HIV, Ebola, many dangerous strains of the Influenza A virus and the coronaviruses mentioned above.

If an outbreak involves a new, unknown pathogen, targeted diagnostic panels are not available at first. Open-view approaches must be used and next-generation sequencing is the method of choice [6, 7]. A swift response is crucial, and analyzing the samples during the sequencing run, as the reads are produced, greatly improves turnaround times. This can be achieved by design using long-read sequencing like Oxford Nanopore (ONT). However, lower throughput and high error rates of those technologies impede their adoption for pathogen detection. Scalability, cost-efficiency and accuracy of Illumina sequencing still make it a gold standard, although it may change in the future with the establishment of improved ONT protocols and computational methods [8].

### Real-time analysis of Illumina sequencing data

Analyzing Illumina reads during the sequencing run poses unique technical and algorithmic challenges. The DRAGEN system relies on field-programmable gate arrays (FPGAs) to speed up the computation and can be combined with specialized protocols to detect clinically relevant variants in the human genome but depends on finished reads [9]. An alternative approach is to use the general purpose computational infrastructure and optimize the algorithms for fast and accurate analysis of incomplete reads as they are produced, during the sequencing run. HiLive [10] and HiLive2 [8] are real-time mappers, performing on par with the traditional mappers like Bowtie2 [11] and BWA [12] with no live-analysis capabilities. However, as read mappers are designed for fast and precise sequence alignment, they are expected to miss most of the reads originating from genomes highly divergent from the available references. Therefore, even though existing live-analysis tools and associated pipelines do cover standard read-based pathogen detection workflows, their performance on novel agents is limited by their dependence on databases of known species. The same problem applies to sequence alignment and taxonomic classification in general, also outside of the real-time analysis context [13]. In this work, we show that using deep learning to predict if a read originates from a human pathogen is a promising alternative to mapping the reads to known references if the correct reference genome is not yet known or unavailable. We also investigate the trade-offs between sequencing time and classification performance.

### Read-based detection of novel pathogens

Deneke et al. [14] have shown that methods like read-mapping [11] (with optional additional filtering steps of PathoScope2 [15]), BLAST [16, 17] or Kraken [18], which all try to assign target sequences to their closest taxonomic matches, fail to yield any predictions for a significant fraction of reads originating from novel pathogens. BLAST was the best of those approaches, missing the least reads and achieving the highest accuracy. More complex detection workflows like PathoScope2 [15], Sigma [19] or KrakenUniq [20] depend on assigning individual reads to taxa by mapping or k-mer matching, so necessarily suffer from the same problems. In contrast, taxonomy-agnostic methods try to reduce their database dependency by assigning putative phenotypes directly to analysed sequences, deliberately omitting the taxonomic classification step. For example, a naïve Bayes classifier based on k-mer frequency features can be trained to classify reads directly into arbitrary classes [21]. However, in the context of detecting novel bacterial pathogens, a random forest approach of PaPrBaG [14] performs much better. Zhang et al. [22] used a kNN classifier to develop a similar method for detection of human-infecting viruses. An analogous deep learning approach, DeePaC, outperforms the traditional machine learning algorithms on both novel bacteria [23] and viruses (DeePaC-vir [24]), offering an additional level of interpretability on nucleotide, read and genome levels. A similar method [25] focuses on detailed predictions for a small set of three viral species and cannot be used in an open-view setting. Preliminary work by Guo et al. [26] supports it, but the code, models or installables are not available yet at the time of writing, so the method cannot be reused. What is more, “novelty” of the viruses in the test set was difficult to assess – it could contain genomes of viruses present in the training set, as long as they were resequenced after 2018. While pathogenicity prediction methods using contigs, whole genomes or protein sets as input also exist, this work focuses on read-based classification to offer real-time predictions and avoid delays necessitated by assembly pipelines. However, read-based methods have been shown to perform well and also for full genomes and assembled contigs, achieving similar or better performance than alignment-based approaches [14, 23, 24]. This mirrors successful adoption of CNNs using raw DNA sequences as inputs in other fields ranging from regulatory genomics [27, 28, 29] to viral bioinformatics [25, 26, 30] and biosecurity [31].

## Methods

### Data preparation

Throughout this paper, we will use the term *subread* in a special sense: the first *k* nucleotides of a given sequencing read (in other words, a *prefix* of a read). The original DeePaC and DeePaC-vir datasets [23, 24] consist of 250bp simulated Illumina reads. The training, validation and held-out test sets contain mixtures of reads originating from different viruses or bacterial species with confirmed labels [32, 33], explicitly modeling generalization to “novel” (i.e. previously unseen) agents (see Supplementary Note 1 for details).

We used the test datasets to generate corresponding subread datasets with subread lengths between 25 and 250 (full read) with a step of 25. Every subread of every subread set had a corresponding subread in all the other sets. Therefore, we explicitly model new information incoming during a sequencing run, as each subread length *k* corresponds to the *k*th cycle. To generalize over a large spectrum of possible subread lengths, we built mixed-length training and validation sets by randomly choosing a different *k* for every read in a set. In this setup, all integer values of *k* between 25 and 250 were allowed. As Bartoszewicz et al. [24] have previously presented both 250bp and 150bp-trained reverse-complement CNN classifier for viruses, we also generated an analogous 150bp subread bacterial dataset, and used it to train a corresponding CNN.

### ResNets and hybrid classifiers

We investigated two architectures shown previously to perform well in the pathogenicity or host-range prediction task – a reverse-complement CNN consisting of 2 convolutional layers and 2 fully-connected layers and a reverse-complement bidirectional LSTM. For more design details and the description of the reverse-complement variants of convolutional and LSTM layers, we refer the reader to [23, 24]. Those architectures guarantee identical predictions for sequences in their forward and reverse-complement orientations in a single forward pass. Previous work [23, 24] has shown that they were more accurate than alternative machine learning and homology-based approaches for the read-based pathogenic or infectious potential prediction task. However, as short subread sequences convey less information, we expected the subread classification problem to be more challenging than in the case of relatively long 250bp reads. We suspected that deeper, more expressive networks could perform better. Therefore, we implemented a new architecture – a reverse-complement ResNet extending the previous work with skip connections [34] while satisfying the reverse-complementarity constraint (see Supplementary Note 2, Table S1 and Fig. S1 for details regarding hyperparameter tuning and selected architectures).

Finally, we create hybrid classifiers, which first extract reads mappable to known references with HiLive2 and then predict the phenotype for the remaining unmapped reads using the ResNet (see Fig. 1 and Supplementary Note 3). This enables identification of the closest relatives of the analyzed pathogen, while still predicting labels for reads missed by the mapper. The reads associated with the pathogenic or infectious phenotype may then be extracted and used in downstream analysis. Note that our workflow, called DeePaC-Live, supports easy substitution of the underlying neural network of any custom Keras model, allowing future improvements in the architecture details and real-time predictions for tasks other than pathogenicity prediction.

**Fig. 1.**
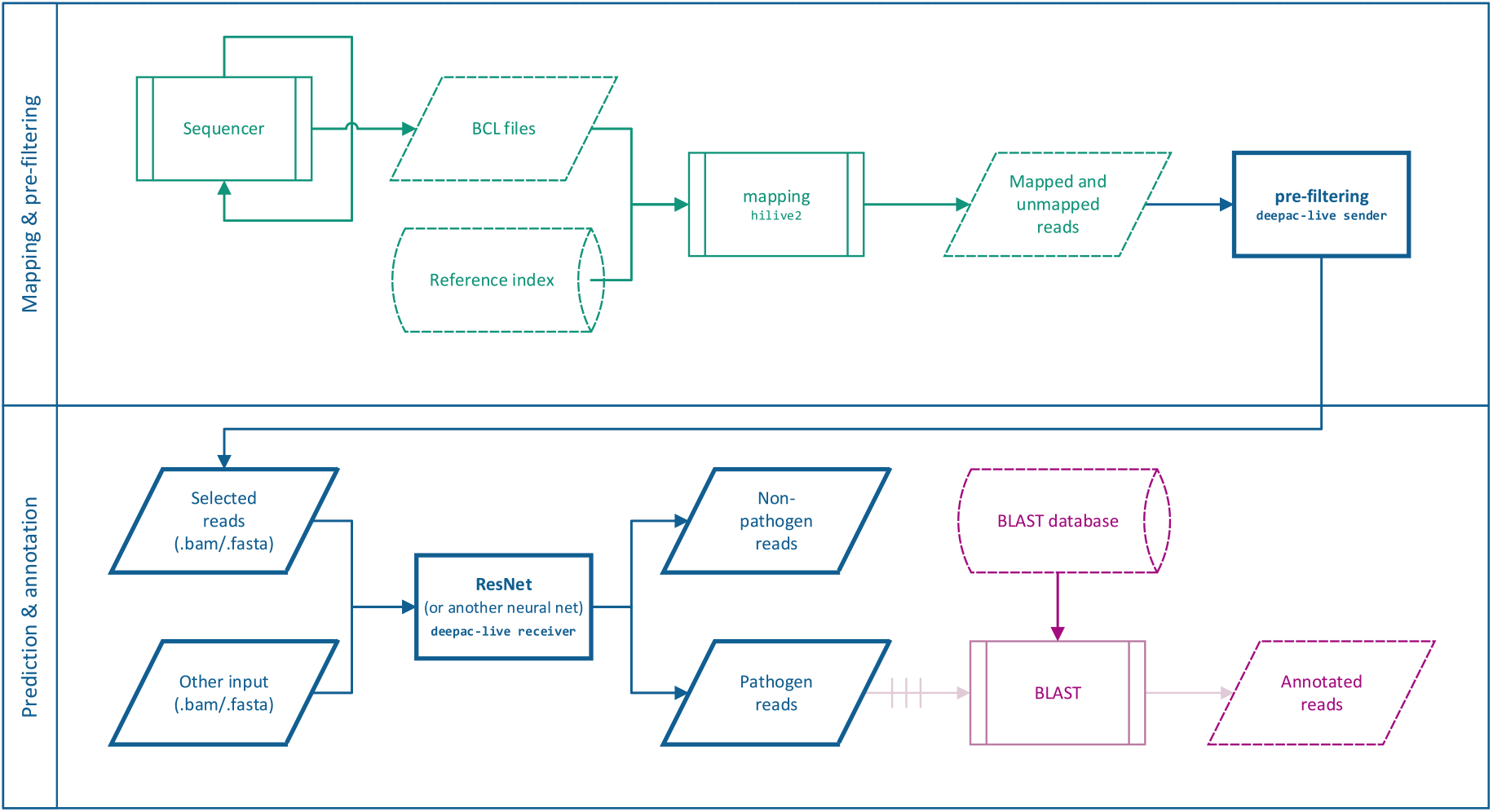
DeePaC-Live workflow, including real-time mapping with HiLive2, classification with ResNets, and optional downstream analysis with BLAST. Mapping and pre-filtering can be performed on the same machine as prediction and annotation, but the pre-filtered reads can also be sent to a GPU-equipped remote server (see Supplementary Note 3 for details regarding integration of HiLive2 and deep learning classifiers). Green: An Illumina sequencer generates the binary base call (BCL) files. They are captured by HiLive2, which maps the reads to an index of known references. Blue: The core of the worklfow, implemented in the DeePaC-Live package. HiLive2 output is captured by the sender module, which selects a subset of reads for further analysis. Selecting the unmapped reads (default) corresponds to the hybrid classifier (HiLive2+ResNet); selecting both mapped and unmapped reads means that all reads will be classified with the ResNet. Alternatively, other inputs in fasta or bam format can be used instead (e.g. basecalled Nanopore reads). The receiver module performs classification of incoming reads with the appropriate ResNet (which can be substituted for another classifier if necessary) and outputs files filtered by the predicted class. Purple: optional follow-up analysis with BLAST, as performed for the Illumina SARS-CoV-2 sequencing run. Putative pathogen reads can be annotated by manually passing them to BLAST using an appropriate database.

### Benchmarking

#### Bacteria

We compare the ResNets and hybrid classifiers to the original DeePaC models, as well as an alternative random forest approach, PaPrBaG [14]. We trained a DNA-only PaPrBaG forest [23] on the mixed-length bacterial dataset. For both machine learning approaches, we average the predictions for both mates of a read pair for a boost in accuracy [23].

In addition to that, we evaluate two alignment-based methods – HiLive2 in the “very-accurate” mode [10, 8] and dc-Megablast [17] with an E-value cutoff of 10 and the default parameters. A successful match to a pathogen reference genome is treated as a positive prediction; a match to a nonpathogen is a negative. In case of multiple matches, the top hit is selected. We build the HiLive2 FM-index and the BLAST database using all the genomes used for training read set generation. If BLAST aligns two mates of a read pair to genomes with conflicting labels (i.e. one pathogen and one non-pathogen), we treat them both as missing predictions. For HiLive2, we treat them separately, as high precision of HiLive2 warrants considering all the obtained matches as relevant. If only one mate has a match, we propagate the match to the other mate. We calculate the performance measures taking all the reads in the sample into account. Hence, missing predictions affect both true positive and true negative rates.

#### Viruses

We use an analogous approach to evaluate the classification performance on reads originating from novel viruses. However, since PaPrBaG is a method developed for bacterial genomes, we benchmark the models against a *k*-nearest neighbors (kNN) virus host classifier [22]. We train the kNN as described by the authors, using non-overlapping 500bp long “contigs” generated from the source genomes. Training based on simulated reads was not possible due to high computational cost, but Zhang et al. [22] showed that a model trained this way can be used to predict pathogenic potentials of short NGS reads. As kNN yields binary predictions, we integrate them using the same approach we use for BLAST. Finally, we compare the models to the original DeePaC-vir models.

#### Real Illumina runs

We evaluate the workflow on real data from *Staphylococcus aureus* (SRA accession: SRR5110368 [35]) and SARS-CoV-2 novel coronavirus (SRR11314339) sequencing runs (see Supplementary Note 4 for preprocessing details). The virus was not present in the training database, as it had not yet been discovered when the DeePaC-vir datasets were compiled. *S. aureus* was also absent from the corresponding training set and was previously used to evaluate DeePaC [23]. To showcase how the approach can be used for rapid detection of novel biological threats, we test the performance of the classifiers after just 50 sequencing cycles. As the predictions of the deep learning approaches do not offer any information about the closest known relative of a novel pathogen, we extend the workflow using BLAST on reads prefiltered by the models. This enables a drastic increase in the pathogen read identification rate while also providing insight into their biological meaning. Using BLAST on full NGS datasets is usually not feasible because of the computational cost. What is more, is has been previously shown [14, 23, 24] that machine learning approaches perform better in pathogenic potential prediction tasks. Therefore, we see the combination of a filtering step with a BLAST follow-up as an in-depth analysis of the subreads of interest while discarding the potentially non-informative ones.

#### Nanopore

Finally, we predict infectious potentials from more noisy subreads of Nanopore long reads. To this end, we resimulated the bacterial and viral datasets using the exact same genomes and the context-independent model of DeepSimulator 1.5 [36]. We set the target average read length to 8kb and discarded reads shorter than 250bp. Then, we extracted 250bp-long subreads for training and evaluation of the classifiers, but kept full reads for benchmarking against minimap2 [37], a popular Nanopore mapper. We chose 250bp as this allows fair comparison with other models and corresponds to information available after ca. 0.5s [38]. Successful predictions after such a short time could be used together with real-time selective sequencing [39] to enrich the samples in reads originating from pathogens and save resources. We trained new models for the bacterial and viral Nanopore datasets and compared them with minimap2 and models trained on 250bp Illumina reads. Evaluation of minimap2 was performed analogously to HiLive2’s, selecting the representative alignment if a chimeric match was found. In addition to the simulated data prepared as explained above, we also used two real SRA datasets: a SARS-CoV-2 isolate (SRR11140745, collected on 14 Feb 2020) and a clinical *S. aureus* sample (SRR8776887 [40]).

## Results

### Subread models

In Illumina paired-end protocols, the barcodes are sequenced after the first mate, making live-demultiplexing possible but problematic. Changing the barcode sequencing order is not trivial, as initial clustering requires sufficient sequence diversity in the first several cycles. A possible workaround uses asynchronous paired-end sequencing protocols [8], sacrificing the first read’s length for faster demultiplexing and relying on the second mate to compensate for the lost information. We tested the models in 100 settings corresponding to different lengths of the first mate (modelling different length-time trade-offs) and the second mate (modelling incoming information after demultiplexing). As shown in the Fig. 2, the previous state-of-the-art for the bacterial dataset [23] is outperformed by the ResNet trained on mixed-length subreads across most of the spectrum of read length combinations. For the longest read pairs, where the sum of read lengths is 400bp or more, accuracy is slightly lower. For the read pairs with the total length below 375bp, the ResNet performs better. This could be related to the old model being explicitly optimized for 250bp-long sequences. Performance of the DeePaC-vir’s 250bp-trained CNN [24] collapses for viral reads shorter than 200bp, while the ResNet trained on the mixed-length reads maintains accuracy higher than 80% for reads as short as 50bp. What is more, the ResNet slightly outperforms the previous state-of-the-art also on full-length reads, with accuracy over 90% for pairs of 225bp or more.

**Fig. 2.**
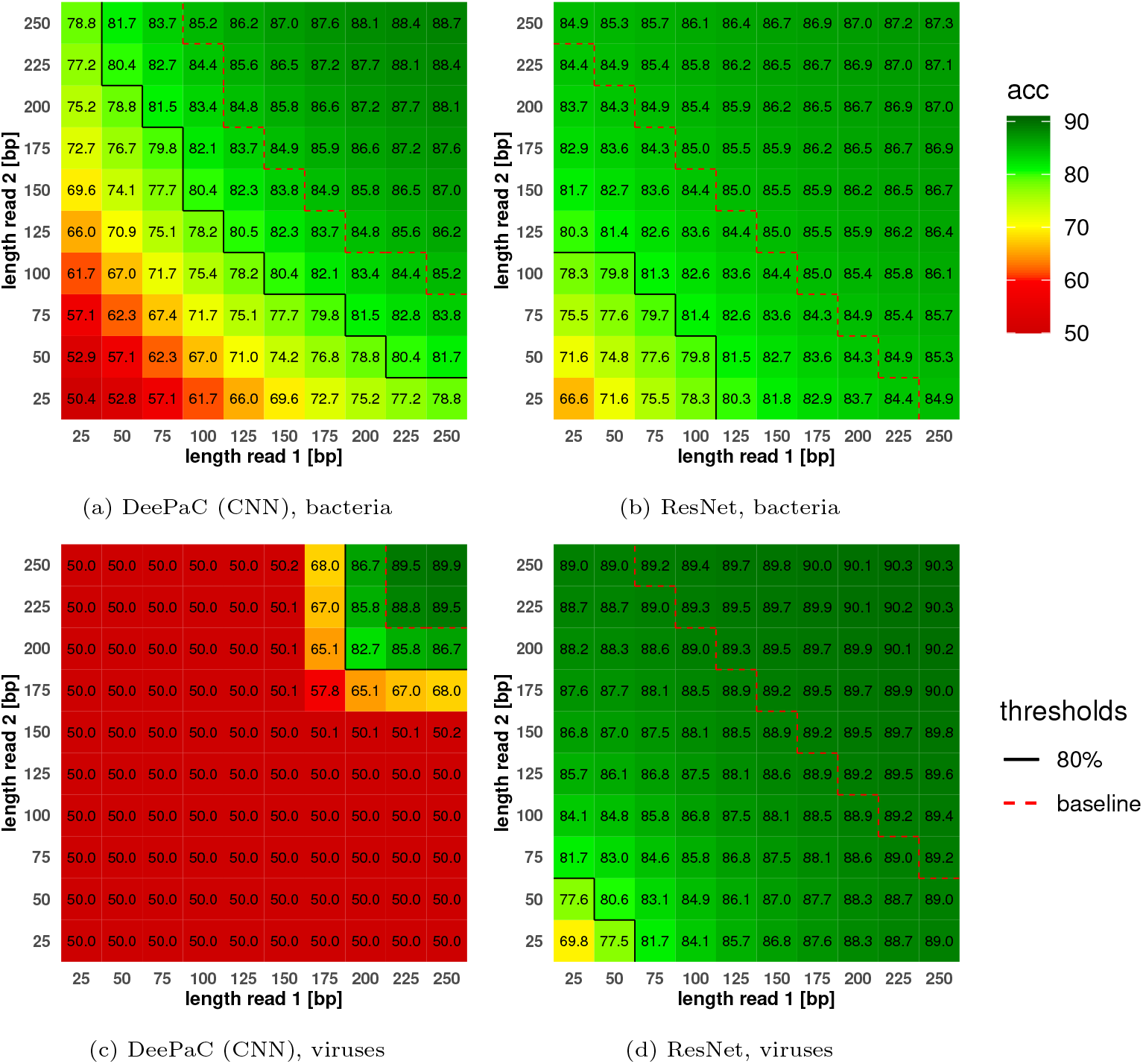
Accuracy heatmaps for different lengths of reads in a read pair. Protocols with asynchronous read lengths (a shorter first mate) allow earlier demultiplexing during the sequencing run. Varying the length of the second mate models obtaining new information during sequencing. The black line corresponds to an arbitrarily set 80% accuracy threshold. The baseline (dashed red line) is the accuracy of each model on 250bp single reads. Crossing the baseline means that the additional sequence information of the second mate contains useful signal, rather than just noise hurting the final predictions. DeePaC (CNN) models were trained on 250bp reads and ResNet on 25-250bp subreads. The accuracy matrices are almost symmetric (with negligible deviations), proving that the models’ performance is on average identical for the first and the second mate.

### Hybrid models

Table 1 and Fig. 3 present classification performance over the whole sequencing run (all cycles for both mates) for the bacterial dataset. The highest accuracy is achieved by the ResNet-based hybrid classifier. High recall of DeePaC (CNN) is actually an artifact – its predictions for shorter subreads are extremely imprecise (precision for 25bp is 50.6%), suggesting that the network simply classifies an overwhelming majority of short subreads as positive regardless of their actual sequence. This effect does not occur for the hybrid classifier, suggesting that although it achieves the second-highest true positive rate overall, it is likely the most sensitive method useful in practice.

**Table 1.** Average performance on reads from novel bacterial species across the whole sequencing run. The hybrid classifier combining HiLive2 and ResNet achieves the highest accuracy. Recall of DeePaC (CNN) is inflated by its unreliable predictions for short subreads. HiLive2 is the most precise method.

|  | Accuracy | Precision | Recall |
| --- | --- | --- | --- |
| HiLive2+ResNet | **83.6** | 80.0 | 89.8 |
| ResNet | 82.1 | 79.2 | 86.7 |
| DeePaC (CNN) | 79.3 | 75.6 | **91.4** |
| DeePaC (LSTM) | 79.7 | 75.2 | 87.0 |
| PaPrBaG | 74.5 | 72.7 | 78.1 |
| BLAST | 60.8 | 84.8 | 76.1 |
| HiLive2 | 22.2 | **97.3** | 36.3 |

**Fig. 3.**
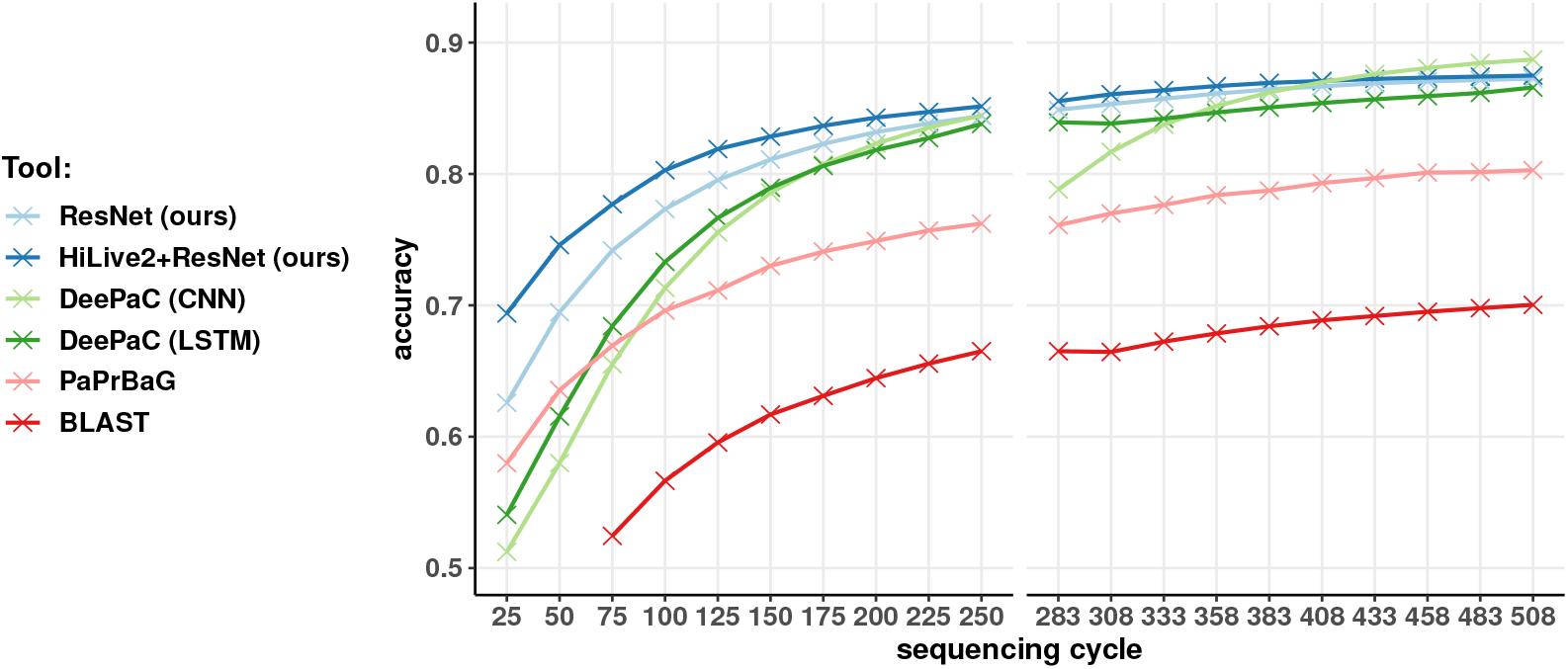
Accuracy for the bacterial dataset. Only the tools achieving more than 60% average accuracy are shown (we omit HiLive2). For BLAST, accuracy values below 50% are not shown. Cycle numbers for the second mate are shifted by 8 due to an 8nt-long simulated barcode present in the BCL files, which is removed at mapping and prediction time. PaPrBaG underperforms even though it was retrained specifically for a subread classification scenario. The mapping approach, represented by HiLive2, is the most precise. Low accuracy of both BLAST and HiLive2 reflects a high missing prediction rate (crossing 80% for HiLive2 at cycles 225-250), although BLAST performs better due to its less strict and more sensitive alignment criteria. Combining HiLive2 with the ResNet results in the best average accuracy overall.

For the viral dataset, the ResNet performs slightly better than HiLive2 even on the reads that HiLive2 is able to map. If only the mapped reads are considered, the ResNet correctly labels 90.7% of them, compared to 89.8% for HiLive2 itself. The effect is especially strong for reads 50bp and longer, where the average accuracy of the ResNet rises to 91.8%, while HiLive2’s stays the same. It seems to stem mainly from increased sensitivity of the deep learning approach (89.3% for reads over 50bp, compared to 81.8% for HiLive) at some cost in precision (95.3% and 99.2%, respectively).

This is most probably why the ResNet has higher accuracy than the hybrid classifier also when unmapped reads are considered, as presented in Table 2 and Fig. 4. It is also the most sensitive prediction method overall. The hybrid classifiers offer a good trade-off between accuracy and precision. As in other real-time analysis tasks [41, 8], the latter is high even for very short subreads (Fig. S2).

**Table 2.** Average performance on reads from novel viruses across the whole sequencing run. ResNet alone achieves the highest accuracy and is the most sensitive method overall, while HiLive2 is the most precise.

|  | Accuracy | Precision | Recall |
| --- | --- | --- | --- |
| HiLive2+ResNet | 85.9 | 93.1 | 77.6 |
| ResNet | **86.5** | 90.5 | **81.5** |
| DeePaC (CNN-150) | 84.8 | 92.3 | 75.6 |
| DeePaC (CNN) | 62.5 | 48.0 | 26.7 |
| DeePaC (LSTM) | 79.0 | 90.0 | 66.0 |
| kNN | 60.7 | 61.7 | 54.1 |
| BLAST | 73.2 | 97.2 | 73.7 |
| HiLive2 | 51.1 | **99.2** | 50.3 |

**Fig. 4.**
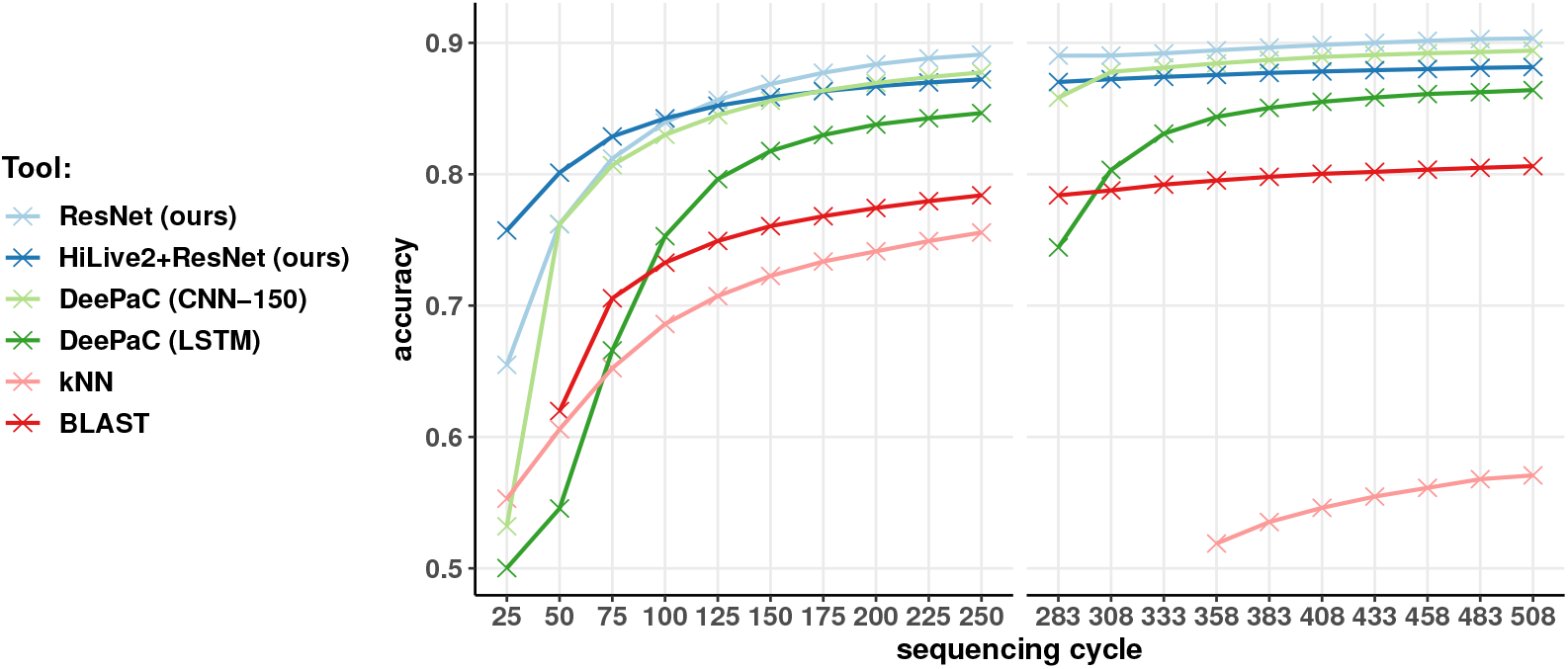
Accuracy for the viral dataset. Only the tools achieving more than 60% average accuracy are shown (we omit HiLive2). For BLAST and kNN, accuracy values below 50% are not shown. DeePaC (CNN-150) corresponds to DeePaC-vir’s CNN_All-150_. DeePaC (CNN) trained on 250bp reads is omitted due to its subpar performance on reads shorter than 200bp (see Table 2 and Fig. 2c). Cycle numbers for the second mate are shifted by 8 due to an 8nt-long simulated barcode present in the BCL files, which is removed at mapping and prediction time. The kNN classifier performs worse than BLAST even for the first mate. Its performance collapses as the second mate is introduced due to many conflicting predictions for both mates, resulting in missing predictions for many read pairs. The ResNet has the best average accuracy overall.

This suggests that the protocol could be useful also beyond the sequencing context, e.g. to improve screening workflows for safe and secure synthetic biology (see Supplementary Note 5), where new methods are needed as engineering of modified pathogens becomes increasingly realistic [42, 43, 44, 45, 46, 47, 48]. Standard screening approaches rely on homology-based pipelines for pathogen detection and functional annotation [48, 49]. Comprehensive evaluation of such systems is challenging, as even attempting to design synthetic DNA encoding novel, harmful functions would not be ethically admissible. Importantly, the screening must work well also for known threats, where evaluation is easier. Assessing performance on novel sequences can only be done indirectly, by removing relevant sequences from the underlying database. This approach has been previously mentioned in [23, 24], where the potential of deep learning as a better alternative to purely homology-based identification was first shown, and is also used here. We note that after the completion of this work, an analogous approach was also presented to test the performance of the independently developed SeqScreen pipeline, explicitly focusing on biosecurity applications [49]. SeqScreen extends homology-based pathogen sequence identification with predicting the functions of proteins to which successful homology hits have been found.

### Runtime

To estimate the sample size that can be analyzed in real-time after parsing or mapping with HiLive2, we measured how many reads per second could be processed by the pathogenicity prediction methods compared in this study. Then, we calculated the number of predictions feasible in a time-frame corresponding to 25 cycles (with wall-time per cycle as in [8]). In the Table S2 we present how many reads can be analyzed with no delays if the output is produced every 25 cycles, together with more detailed information on the hardware used and the effect of adjusting the inference batch size. The 18-layer ResNet is faster than the original DeePaC models and only marginally slower than optimized CNNs and LSTMs, analyzing over 56 million reads in the given time-frame (5656 reads/s). This is over 6 times faster than the best non-deep learning based method for both bacteria and viruses, at a much lower cost in terms of the computational resources. This prediction speed is enough to guarantee real-time predictions for Illumina iSeq 100, MiniSeq and MiSeq devices, with a maximum of 4 and 25 million reads per run. For sequencers with even higher maximum throughput, like the obsolesced HiSeq and newer NovaSeq 550 machines (with up to 400 million reads per run), further speed-up is possible by distributing the computation across more than one GPU. Multiple receiver instances can be easily assigned to dedicated GPUs to handle different barcodes, cycles or both in parallel. ResNet training time for the maximum of 30 epochs corresponded to 48h on two Tesla V100 GPUs; trained models are available for future use (see “Data availability”).

### Real sequencing data

We benchmarked the best bacterial model on data from a real *S. aureus* sequencing run. As this dataset contains reads from just one “novel” pathogen (a species which was not present in the training database), the true positive rate (recall) and accuracy are equivalent. The hybrid HiLive2+ResNet classifier crosses the 90% threshold just after 75 cycles (when the true positive rate equals 89.8%) and reaches 98.8% in the last analyzed cycle (Fig. S3). HiLive2 is only able to identify 5.8% of the reads at its best cycle, which drops down to 2.0% at the end of the sequencing run, when longer sequences are analyzed.

We further evaluated the approach on data from a real SARS-CoV-2 sequencing run. Note that the training database did not contain a SARS-CoV-2 reference genome, mimicking the pre-pandemic state of knowledge. In this setting, we used BLAST as an example follow-up analysis of the reads filtered with the hybrid classifier or the pure deep learning approach after just 50 cycles (Table 3). As in the case of the open-view DeePaC-vir dataset, the neural network itself performs better than the hybrid classifier, being more accurate even on the reads mappable with HiLive2. More specifically, HiLive2 suffers from a high false negative rate of 96.3% even when only the mapped reads are considered, which is probably because it was designed for mapping against known references. The ResNet is substantially more accurate on the same reads (with a recall of 10.9%, compared to HiLive2’s 0.6%). Because of that, omitting the mapping functionality of HiLive2 (and using it just for parsing the BCL files generated by the sequencer) results in better performance. Notably, even the spurious non-pathogenic identifications of HiLive2 can be useful – 99.7% of the mapped reads are identified as originating from coronaviruses, and 90.5% are identified as bat coronaviruses, including the *Rhinolophus* (horseshoe bat) coronaviruses and bat SARS-like viruses, which are probably closely related to SARS-CoV-2.

**Table 3.** Reads identified as pathogenic from the SARS-CoV-2 sequencing run. The ResNet alone is able to identify the most reads, but cannot annotate them with matches to the closest known references. It is more accurate than HiLive2 even on the reads that the latter is able to map, which explains why it performs better than the HiLive2+ResNet hybrid classifier. Combining HiLive2 or BLAST with the ResNet identifies taxonomic signals while extracting more reads than pure mapping. BLAST output can be used to annotate the reads with the closest taxonomic match only (annot.), or to form a consensus predictor (cons.) by selecting subreads assigned to the pathogenic class by both the ResNet and BLAST.

|  | Recall | Annot. rate |
| --- | --- | --- |
| HiLive2 | 0.6% | 0.6% |
| HiLive2+ResNet | 41.0% | 0.6% |
| ResNet | **51.3%** | 0.0% |
| HiLive2+ResNet+BLAST (annot.) | 41.0% | 27.7% |
| ResNet+BLAST (annot.) | **51.3%** | **37.9%** |
| HiLive2+ResNet+BLAST (cons.) | 6.8% | 6.8% |
| ResNet+BLAST (cons.) | 9.3% | 9.3% |

Similar identifications can be made with BLAST on the much larger set of ResNet-filtered subreads. As the deep learning models consistently outperform BLAST, we use them as predictors to extract subreads of interest. A BLAST follow-up analysis annotates the selected 50bp subreads with their closest taxonomic matches (which may include non-pathogens) wherever a match is found. Alternatively, we can create a consensus predictor by treating BLAST as a confirmatory analysis to focus only on subreads which are predicted to originate from the positive class *and* have a positive BLAST match. In the latter case, we observe a significant enrichment in sequences more similar to the pathogenic SARS-CoV-1 virus. 99.3% of the subreads identified as “pathogenic” by the consensus ResNet+BLAST workflow are matched with human SARS viruses present in the training database, while the number of identified subreads is almost 15 times higher than for HiLive2 alone. The results suggest that predictions of the ResNet, even without a BLAST follow-up, are also reliable, while offering a recall rate 80 times higher than HiLive2. However, further analysis steps (with BLAST or other approaches, e.g. taxonomic classifiers) are required to gain more fine-grained insights into the origin of ResNet-filtered reads.

### Nanopore reads

Finally, we evaluated the Nanopore models to investigate possible applications to noisier long-read sequencing technologies (Table 4). The Nanopore-trained ResNets achieved higher validation accuracy than the CNNs and LSTMs trained on the same data and were selected for further evaluation. As expected, mapping with minimap2 is the most precise method and Illumina-trained neural networks of DeePaC underperform in this context. Noisy reads are especially challenging for Illumina-trained LSTMs. Their precision and true positive rates become unstable, resulting in relatively low accuracy. Using Nanopore error models for training promotes more robust models. Strikingly, first 250bp of a read are enough for the ResNets to noticeably outperform minimap2, even when it uses whole reads. This holds for real data as well. When the correct reference is not yet available (as before the pandemic), minimap2 recalls 66.9% of full-length *S. aureus* reads and only 9.9% of full SARS-CoV-2 reads, compared to 94.7% and 52.7% for the ResNets respectively (Table S3).

**Table 4.** Performance on Nanopore data. Minimap2 was evaluated on both full reads and 250bp subreads. ResNets were trained on Nanopore data with identical species composition as the Illumina data used for DeePaC CNNs and LSTMs. and evaluated on 250bp subreads. Minimap2 yields no matches for between 13% (viruses, full length) and 69% (bacteria, 250bp) of the reads. Acc. – accuracy, Prec. – precision, Rec. – recall.

|  | Bacteria |  |  | Viruses |  |  |
| --- | --- | --- | --- | --- | --- | --- |
|  | Acc. | Prec. | Rec. | Acc. | Prec. | Rec. |
| ResNet | **78.5** | 73.4 | 89.3 | **88.5** | 90.3 | **86.4** |
| DeePaC (CNN) | 77.8 | 73.1 | 87.8 | 81.4 | 84.0 | 77.6 |
| DeePaC (LSTM) | 73.7 | 66.5 | **95.7** | 78.5 | 92.3 | 62.1 |
| minimap2 (250bp) | 30.5 | **97.2** | 46.6 | 59.3 | **99.2** | 61.8 |
| minimap2 (full) | 66.7 | 91.4 | 79.7 | 80.2 | 98.9 | 82.4 |

These results suggest that the classifiers can find applications in selective sequencing workflows, enabling targeted analysis of reads originating from novel pathogens while discarding potentially less-interesting non-pathogen reads. Although a given read could contain sequences matching to pathogen references located after the initial 250bp, the risk of premature termination seems to be mitigated by the classifier’s superior performance, especially in the case of novel viruses. This risk can be further adjusted to the user’s needs by selecting an alternative classification threshold, manipulating the expected sensitivity, precision and false positive rates as shown by the ROC and PR curves (Fig. S4).

## Discussion

All the limitations of the previously described read-based methods of pathogenic potential prediction [14, 22, 23, 24] apply to this study too. The models presented here assign probability-like scores to DNA and RNA sequences without establishing a mechanistic link between a given sequence and the predicted phenotype. This is, however, an advantage of the proposed approach as well. By resigning from speculation on that link (e.g. by sequence alignment to known relatives), models trained on carefully selected data outperform the traditional methods in terms of both prediction speed and accuracy, yielding predictions for all sequences in the sample. Any assumptions and biases affecting the labels will be reflected by the trained classifier. While this is also the case for read alignment and k-mer based classification, they do not require retraining after a database update. On the other hand, as the classifiers generalize well to sequences absent from the training database, they may be updated less frequently while maintaining the desired performance.

A separate question is whether the captured signal has any underlying functional meaning, or is purely taxonomic in nature. BLAST, as a sensitive method of homology detection, is a gold-standard for finding taxonomic relationships. Therefore, it could be assumed that outperforming BLAST is a sign of learning more than just evolutionary distances. On the other hand, the very nature of read-based predictions renders any broader biological context inaccessible – the analyzed sequences are simply too short to contain reliable peptide-level features [14, 23], let alone information about the structural or functional characteristics of the encoded proteins (or intergenic regions). However, intepretability workflows like Genome-Wide Phenotype Potential Analysis [24] have shown that read-based pathogenicity prediction models assign high pathogenic potentials to reads originating from genes engaged in virulence, also specifically in the case of *S. aureus*. What is more, they show that regions of higher pathogenic potential are non-uniformly distributed in both bacterial and viral genomes with “peaks” of elevated potential aligning with relevant genes. This suggests that even though detecting pathogenicity islands or virulence factors directly is not possible, reads associated with the pathogenic phenotype do originate from important regions of interest. On the other hand, one can expect similar sequences to yield similar predictions. This makes distinguishing between closely related pathogens and non-pathogens challenging, as shown previously for SARS-CoV-2 and its relative, RaTG13 [24]. The problem can be explicitly modelled by including related viruses infecting different hosts (as in the DeePaC-vir dataset used in this study) or by training classifiers targeting novel strains of known bacterial species [23]. Nevertheless, occasional misclassifications of similar sequences are a real possibility that has to be kept in mind. This also applies to alignment-based and k-mer based approaches.

The accuracy achieved by the models clearly shows that predicting if a read comes from a bacterial pathogen or a human-infecting virus is indeed possible, even if there is no reference genome available. This may actually be a form of texture bias – CNNs have been shown to correctly classify images based on fragments of as little as 17×17 pixels [50]. Here, a local DNA or RNA pattern is often predictive of the phenotype label assigned to the genome. However, the models do not return any information on the closest possible match, which is generally necessary in any pathogen detection task. In this study, we proposed to solve this problem by combining the deep learning approach with an alignment-based one. Alternative methods of taxonomic classification could be used instead – based on either k-mers or machine learning. Kraken [18], a k-mer approach, was outperformed by both BLAST and PaPrBaG [14]. Nevertheless, we can imagine that using a well-trained taxonomic classifier, preferably one yielding at least putative species-level predictions for every read, would be a very useful tool for follow-up analyses of reads prefiltered with the models. On the other hand, since the reads associated with pathogenicity are often co-localized within relevant genomic features [24], assembly of the filtered reads could recover longer contigs corresponding to genes or perhaps even gene clusters.

A combination of mapping and ResNets could also form a part of more complex real-time pathogen detection workflows like PathoLive [51]. For example, PAIPline [52] identifies pathogens in metagenomic and clinical samples via mapping and a BLAST follow-up analysis, but can only start the analysis after the sequencing is finished. Exchanging Bowtie2 [11] for HiLive2 with a hybrid classifier directing potentially informative reads to the BLAST confirmatory step could serve as a backbone of an extended, real-time version of the pipeline. Alternatively, PathoLive could be extended with ResNet and BLAST follow-up steps. To fully handle metagenomic samples, the classifiers would have to be retrained in a multi-class setting encompassing a broader spectrum of clinically relevant pathogen groups. On the other hand, if there are reasons to believe that the disease-causing agent is a virus or a bacterium, the models presented here may suffice.

As our classifiers rely on either BAM or fasta input, they are not necessarily dependent on HiLive2 and can be used in combination with alternative approaches to accelerated sequencing analysis, for example the DRAGEN system. Nanopore-trained models perform relatively well despite higher sequencing noise, and we imagine incorporating pathogenicity prediction into real-time selective sequencing workflows [39]. Since 250bp subreads are enough to make predictions more accurate than possible with mapping even fully sequenced reads, it would be possible to terminate sequencing of some reads quickly to focus on sequencing those originating from pathogens.

## Key points

- We present a new workflow for real-time prediction of pathogenic potential of novel bacteria and viruses, accessing the intermediate files of an Illumina sequencer.
- Deep learning models specialized in inference from incomplete short- and long-read sequencing data outperform alternatives on both simulated and real reads.
- Combining deep learning with homology search recovers reads originating from novel pathogens without sacrificing performance on known agents, highlighting close relatives.
- The protocol could also be used for sequence-based tasks beyond NGS analysis, for example as a screening system for synthetic DNA sequences difficult to evaluate before.

## Data availability

The workflow is available as a command-line tool and a Python package, DeePaC-Live. It can be installed with Bioconda [53], Docker or pip. The code and installation instructions are available at https://gitlab.com/dacs-hpi/deepac-live (real-time inference and HiLive2 integration) and https://gitlab.com/dacs-hpi/deepac (ResNet training and data preprocessing). The datasets are hosted at https://doi.org/10.5281/zenodo.4456857 along the trained models and config files describing the model architecture and hyperparameters (https://doi.org/10.5281/zenodo.4456008

## Competing interests

There is NO Competing Interest.

## Author contributions statement

J.M.B. and B.Y.R. conceived the study, J.M.B. and U.G. conducted the experiments and analysed the results. J.M.B. wrote the manuscript. All authors read, revised and approved the manuscript.

## Acknowledgments

We thank Tobias P. Loka and Melania Nowicka for multiple valuable discussions and comments. This work was supported by the German Academic Scholarship Foundation (J.M.B.), the BMBF-funded Computational Life Science initiative (project DeepPath, 031L0208, to B.Y.R.) and the BMBF-funded de.NBI Cloud within the German Network for Bioinformatics Infrastructure (de.NBI) (031A537B, 031A533A, 031A538A, 031A533B, 031A535A, 031A537C, 031A534A, 031A532B).

**Jakub M. Bartoszewicz** is a PhD candidate at Free University Berlin, a research assistant at the Hasso Plattner Institute and a visiting scientist at the Robert Koch Institute.

**Ulrich Genske** is a research assistant at Charité – Universitätsmedizin Berlin and Hasso Plattner Institute. He was previously a research assistant at the Robert Koch Institute.

**Bernhard Y. Renard** is professor for Data Analytics and Computational Statistics at Hasso Plattner Institute, Digital Engineering Faculty of the University of Potsdam. Earlier, he built up the bioinformatics division at Robert Koch Institute.

## Notes

### Competing Interest Statement

The authors have declared no competing interest.

