## Supplementary Information for "Deep learning-based real-time detection of novel pathogens during sequencing"

Jakub M. Bartoszewicz [1-3], Ulrich Genske [1-4] and Bernhard Y. Renard [1-2]

### Supplementary Notes

#### Supplementary Note 1: DeePaC datasets

The original DeePaC dataset consists of 250bp simulated Illumina reads in fastq format. The training, validation and test sets contain reads originating from different species of pathogenic (including opportunistic pathogens) or commensal bacteria labelled using the IMG database [Chen et al., 2019]. The DeePaC-vir dataset is built in an analogous way using different viruses mined from the Virus-Host Database [Mihara et al., 2016]. Three alternative versions of the viral dataset are available, differing in the negative class definition. We used the fully open-view "All" dataset, containing all viruses available in VHDB. In all cases, the training set contained 20 million single reads, the validation set contained 2.5 million single reads and the test set – 2.5 million paired-end reads. This setup allows training models correctly handling single, isolated reads, but also testing their performance on read pairs. All sets were balanced with regard to the class distribution and contained a mixture of reads originating from multiple different species. Most importantly, the training, validation and held-out test sets contain different viruses or bacterial species, so that generalization to "novel" agents (i.e. unseen in training) can be explicitly evaluated. For more details regarding the dataset generation, we refer the reader to the corresponding publications [Bartoszewicz et al., 2020, 2021].

#### Supplementary Note 2: Hyperparameter tuning

We considered 18- and 34-layer ResNet variants where all convolutional layers of a standard ResNet (including size-1 convolutions in skip connections) are replaced with reverse-complement convolutions [Bartoszewicz et al., 2020]. We trained them for a maximum of 30 epochs, using early stopping with a patience of 10 epochs (see Table S2 and Fig. S1 for architecture details). For all models, we used input dropout, which may be understood as switching a random fraction of the input nucleotides to $N$s. As generating subreads already discards some sequence information, we retuned the input dropout rate for the bacterial models, testing the values of 0.2 and 0.25. For the viral models, it was already shown that the dropout rate of 0.25 works better even in the case of 150bp subreads; we therefore only considered the higher value. We compared the CNN, LSTM and ResNet models trained on the mixed-length datasets; in addition to that we also considered the bacterial CNN trained on 150bp subreads analogous to the viral $\mathrm{CNN}_{\mathrm{All\text{-}150}}$ from Bartoszewicz et al. [2021]. The ResNet-18 trained with an input dropout rate of 0.25 achieved the highest accuracy on the bacterial mixed-length validation set and was selected for further evaluation. For viruses, the ResNet models were the best as well – although the ResNet-34 was the most accurate in absolute terms, the error rate improvement over the 18-layer variant was negligible (<0.5%) while the computational cost (measured in wall-clock time of both training and inference) was roughly twice as high. Since inference speed is crucial for the

[1]Hasso Plattner Institute, Digital Engineering Faculty, University of Postdam, Potsdam, Brandenburg, Germany
[2]Bioinformatics (MF1), Department of Methodology and Research Infrastructure, Robert Koch Institute, Berlin, Germany
[3]Department of Mathematics and Computer Science, Free University of Berlin, Berlin, Germany
[4]Department of Radiology, Charité – Universitätsmedizin Berlin, Free University of Berlin, Humboldt University, and Berlin Institute of Health, Berlin, Germany

application presented here, we decided to select the equally accurate but faster and more efficient ResNet-18. For Nanopore data, we retrained only the CNNs, LSTMs and ResNets-18, omitting the ResNet-34 architectures.

#### Supplementary Note 3: HiLive2 and ResNet integration

To capture Illumina reads as they are generated by a sequencer, we use HiLive2's BCL file conversion and real-time mapping capabilities [Lindner et al., 2017, Loka et al., 2019]. The tool, DeePaC-Live, consists of three asynchronously callable modules. The sender module watches the HiLive2 output directory, detecting BAM files with both mapped and unmapped reads. By default, it selects only the unmapped reads for further analysis, but this can be adjusted by the user to focus on either mapped reads, or all sequenced reads. The output of the sender module may be automatically sent over to a remote server (e.g. a GPU-equipped machine) using the SFTP protocol. Data privacy issues should be kept in mind.

The receiver module may operate on the remote or local machine depending on the available infrastructure. It captures the sender's output and uses a selected deep neural network to predict pathogenic potentials (standard sigmoid output scores between 0 and 1) for all the selected reads. Then, it filters them according to a predefined decision threshold (typically 0.5), outputting separate files for reads associated with a pathogenic and nonpathogenic phenotype. Finally, the optional refiltering module allows reanalyzing the prediction with an alternative threshold (e.g. to select only the highest-confidence predictions) and averaging the outputs of multiple receiver modules to create a simple ensemble classifier.

Any custom Keras model can be used for predictions, including arbitrary binary classifiers for tasks other than described here. We also support seamless integration with the built-in DeePaC [Bartoszewicz et al., 2020] and DeePaC-vir [Bartoszewicz et al., 2021] models. However, previously available models were optimized for a relatively long read length of 250bp (with one viral model trained for 150bp reads). We suspected that they would underperform in a real-time analysis scenario, where much shorter reads are analyzed. Therefore, we trained new models, aiming to achieve high performance for both the intermediate cycles and the final output of the sequencer. As the prediction functions of DeePaC were not optimized for fast inference, we added a possibility to adjust the inference batch size to fully utilize computing power of a given GPU. We set the batch size to 1536, being the highest multiple of 512 that would not cause out-of-memory errors for any of the tested models. Note that while the batch size could be further increased for the CNN and ResNet-based networks used in this study, this did not speed up inference any more.

#### Supplementary Note 4: Real data preprocessing details

To test the performance of the bacterial models on real sequencing data, we analyze reads coming from a real sequencing run of a pathogenic bacterium *Staphylococcus aureus*. This species was not present in the training set (it had been randomly placed in the validation set), so models a "novel" pathogen without a known reference genome. The same species was used previously by Bartoszewicz et al. [2020] to assess the original version of DeePaC; here, we focus on analyzing the sequences as they are generated by the sequencer as opposed to predicting for full reads after the sequencing run is finished. To this end, we downloaded an SRA archive of 251bp-long paired-end reads (accession number SRR5110368) sequenced with an Illumina MiSeq device [Manara et al., 2018]. To evaluate the viral models, we downloaded an archive of 151bp-long paired-end SARS-CoV-2 reads originating from a COVID-19 positive human from San Diego county (SRR11314339). We use untrimmed reads with the quality information to generate BCL files as they would be internally generated by the sequencer. We then run HiLive2 on the BCL data to map the reads to the training reference database; HiLive2 output is then parsed and passed to the models. However, we ignore the last cycle of each mate when generating HiLive2 output and subsequent analyses, as the bad quality of this last nucleotide makes it generally unreliable. We select the predictor that achieves the highest average accuracy on the DeePaC or DeePaC-vir dataset and compare it to the standard, mapping-based real-time analysis with HiLive2 alone.

#### Supplementary Note 5: Biosecurity and biosafety in synthetic biology

Methods developed for pathogen detection and real-time sequencing could also be used beyond the sequencing context. For example, the models presented here could be used at the DNA synthesis facilities to screen ordered sequences against potential biosecurity threats. Host-range of viral pathogens can be deliberately modified [Herfst et al., 2012, Imai et al., 2012], and a virus similar to the Variola virus (the cause of smallpox and a bioweapon) was synthesized [Noyce et al., 2018, Thiel, 2018]. Lipsitch and Inglesby [2014] speculated on modifications increasing the pathogenicity of coronaviruses. On the other hand, a report by the National Academies of Sciences, Engineering, and Medicine [2018] sees virulence-enhancing manipulation of existing bacteria as the issue of the highest concern. Computational screening of ordered sequences is a standard, but challenging precaution measure used by the DNA synthesis industry; evaluation of novel sequences requires significant computational resources and expert analysts. As it depends on sequence alignment against databases of known threats, it suffers from the same problems as other taxonomy-dependent pathogen detection methods. Evaluating sequences shorter than 200bp is usually not feasible due to high false positive rates and the computational burden; a PhD-level proficiency in bioinformatics is required to both implement the pipelines and analyse the results [Diggans and Leproust, 2019]. Taken together, those challenges warrant investigating deep learning alternatives to traditional workflows. The models deliver high accuracy and precision for sequences well below the established 200bp limit, and their false positive rates can be lowered even more if a decision threshold higher than the default 0.5 is used. Given the inference speed of the classifiers, we envision a system where the suspicious sequences are filtered with DeePaC-Live and piped into a follow-up analysis akin to BLAST, lowering the computational burden of sequence alignment and improving the performance.

### Supplementary Tables and Figures

Table S1: A summary of the datasets used in this study. Simulated datasets have been prepared based on datasets of Bartoszewicz et al. [2020, 2021] (see Supplementary Note 1). Subread test sets with read lengths between 25 and 250 (step of 25) were generated based on the original simulated Illumina test sets; we only list the them once in the table for clarity.

| content | technology | positive reads | negative reads | accession number |
|---|---|---|---|---|
| Bacteria (train.) | Illumina (sim.) | 10M | 10M | https://zenodo.org/record/4456857 |
| Bacteria (val.) | Illumina (sim.) | 1.25M | 1.25M | https://zenodo.org/record/4456857 |
| Bacteria (test) | Illumina (sim.) | 2x0.625M | 2x0.625M | https://zenodo.org/record/3678563 |
| Viruses (train.) | Illumina (sim.) | 10M | 10M | https://zenodo.org/record/4456857 |
| Viruses (val.) | Illumina (sim.) | 1.25M | 1.25M | https://zenodo.org/record/4456857 |
| Viruses (test) | Illumina (sim.) | 2x0.625M | 2x0.625M | https://zenodo.org/record/4312525 |
| Bacteria (train.) | Nanopore (sim.) | 10M | 10M | https://zenodo.org/record/4456857 |
| Bacteria (val.) | Nanopore (sim.) | 1.25M | 1.25M | https://zenodo.org/record/4456857 |
| Bacteria (test) | Nanopore (sim.) | 1.25M | 1.25M | https://zenodo.org/record/4456857 |
| Viruses (train.) | Nanopore (sim.) | 10M | 10M | https://zenodo.org/record/4456857 |
| Viruses (val.) | Nanopore (sim.) | 1.25M | 1.25M | https://zenodo.org/record/4456857 |
| Viruses (test) | Nanopore (sim.) | 1.25M | 1.25M | https://zenodo.org/record/4456857 |
| *S. aureus* | Illumina | 2x1.1M | 0 | SRR5110368 |
| SARS-CoV-2 | Illumina | 2x517.3k | 0 | SRR11314339 |
| *S. aureus* | Nanopore | 83.4k | 0 | SRR8776887 |
| SARS-CoV-2 | Nanopore | 396.4k | 0 | SRR11140745 |

Table S2: ResNet architecture details. Conv1 and first layers of stages conv3-conv5 use a stride of 2, and all other layer use the stride of 1. Stages 2-5 consist of multiple layers with the same filter width and number of filters. Batch normalization is used after all hidden layers. After the convolutions, we use global average pooling and a fully-connected output layer.

| stage | ResNet-18 | ResNet-34 |
|---|---|---|
| conv1 | filter width:7, filters:64 | filter width:7, filters:64 |
| conv2 | [filter width:5, filters:64] x 4 | [filter width:5, filters:64] x 6 |
| conv3 | [filter width:5, filters:128] x 4 | [filter width:5, filters:128] x 8 |
| conv4 | [filter width:5, filters:256] x 4 | [filter width:5, filters:256] x 12 |
| conv5 | [filter width:5, filters:512] x 4 | [filter width:5, filters:512] x 6 |
| pool | global average pooling | global average pooling |
| out | 1-unit fully-connected | 1-unit fully-connected |

Table S3: Inference speed in reads per second and reads per sequencing time of 25 cycles. This is an inherently difficult comparison, as inference with neural networks can be accelerated with GPUs, while other methods cannot. We used a desktop computer equipped with a consumer-grade GPU to benchmark the throughput of deep learning approaches, and a 128-core machine with 500 GB RAM for the other methods. All runtimes were calculated for single 250bp reads. Running PaPrBaG on more than 8 cores was not possible due to high memory consumption (over 400GiB, compared to less than 4GiB for the CNNs, LSTMs and ResNets). BLAST database size influences alignment time. Therefore, we present separate runtimes for the bacterial ($BLAST_{Bac}$) and viral ($BLAST_{Vir}$) dataset. Top four models (bold) offer relatively similar performance, crossing the arbitrary threshold of 5000 classified reads per second on a single consumer-grade GPU. CNN (adj.) and LSTM (adj.) differ from the previously published DeePaC versions only by the adjusted inference batch size (see Supplementary Note 3).

| | Device | Reads/s | Reads/25c |
|---|---|---|---|
| **CNN (adj.)** | 1x Nvidia RTX 2080 Ti | **6313** | **63.1M** |
| **LSTM (adj.)** | 1x Nvidia RTX 2080 Ti | **5896** | **58.9M** |
| **ResNet** | 1x Nvidia RTX 2080 Ti | **5656** | **56.5M** |
| **DeePaC (CNN)** | 1x Nvidia RTX 2080 Ti | **5000** | **50.0M** |
| ResNet-34 | 1x Nvidia RTX 2080 Ti | 2880 | 28.8M |
| DeePaC (LSTM) | 1x Nvidia RTX 2080 Ti | 1855 | 18.5M |
| PaPrBaG | 8x Intel Xeon E5-4667 v4 @ 2.20Ghz | 906 | 9.0M |
| $BLAST_{Vir}$ | 100x Intel Xeon E5-4667 v4 @ 2.20Ghz | 833 | 8.3M |
| kNN | 100x Intel Xeon E5-4667 v4 @ 2.20Ghz | 37 | 0.3M |
| $BLAST_{Bac}$ | 100x Intel Xeon E5-4667 v4 @ 2.20Ghz | 160 | 1.6M |

Table S4: Performance on real Nanopore data. Minimap2 was evaluated on both full reads and 250bp subreads. ResNets were trained on Nanopore data with identical species composition as the Illumina data used for DeePaC CNNs and LSTMs. and evaluated on 250bp subreads. Mean sequencing time and maximum sequencing time per read correspond to estimated sequencing times for the mean read length and maximum read length in a dataset (assuming the sequencing speed of 500bp/s).

| | *S. aureus* (SRR8776887) | | | SARS-CoV-2 (SRR11140745) | | |
|---|---|---|---|---|---|---|
| | Recall | Mean seq. time | Max. seq. time | Recall | Mean seq. time | Max. seq. time |
| ResNet | **94.7** | 0.5 s/read | 0.5 s/read | **52.7** | 0.5 s/read | 0.5 s/read |
| minimap2 (250bp) | 3.3 | 0.5 s/read | 0.5 s/read | 4.6 | 0.5 s/read | 0.5 s/read |
| minimap2 (full) | 66.9 | 15.9 s/read | 107.2 s/read | 9.9 | 1.3 s/read | 12.7 s/read |

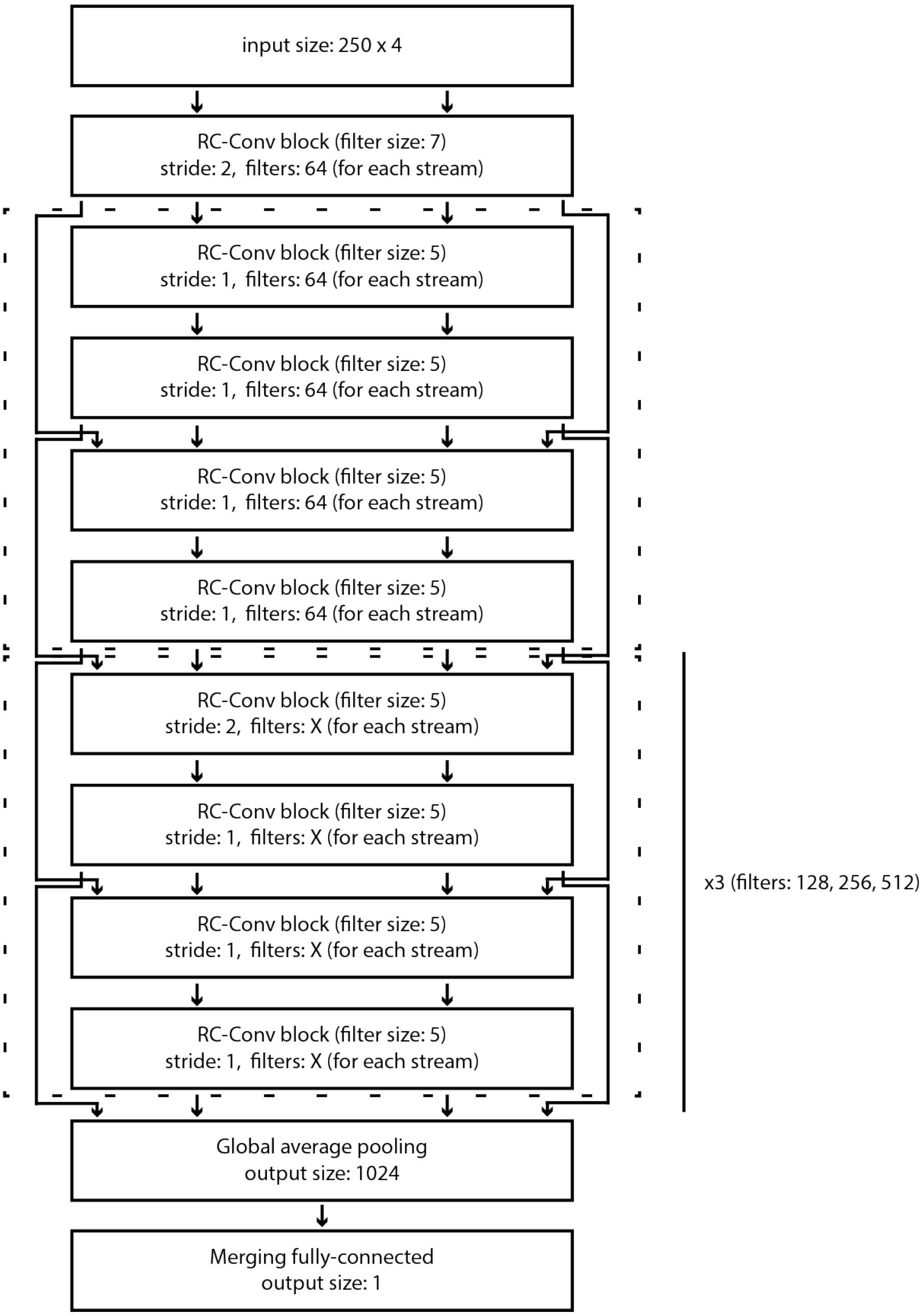


Figure S1: A simplified visualization of the 18-layer variant of the reverse-complement ResNet used in this study (see Table S1). The reverse-complement convolution blocks (RC-Conv) output feature maps in the forward and reverse-complement orientations, represented as small arrows between blocks (the "forward" and "reverse-complement" streams). The skip connections (larger arrows) maintain invariance to reverse-complementarity by either simply summing the outputs of two layers (as in a standard ResNet) or by applying a size-1 RC-convolution if the layer dimensions do not match (similarly to how a standard ResNet uses standard size-1 convolutions). Note that in contrast to a standard ResNet, all convolutional layers are 1D-convolutions, since the inputs are one-hot encoded nucleotide sequences. The first dashed rectangle corresponds to the conv2 stage in Table S1, and the second dashed rectangle represents stages conv3-conv5. The architecture presented here can be reproduced using the `deepac train -c config.ini` command, where `config.ini` is one of the config files available at https://doi.org/10.5281/zenodo.4456008 along the trained models. `deepac` is installed automatically with `deepac-live`. The necessary training and validation datasets are hosted at https://doi.org/10.5281/zenodo.4456857.

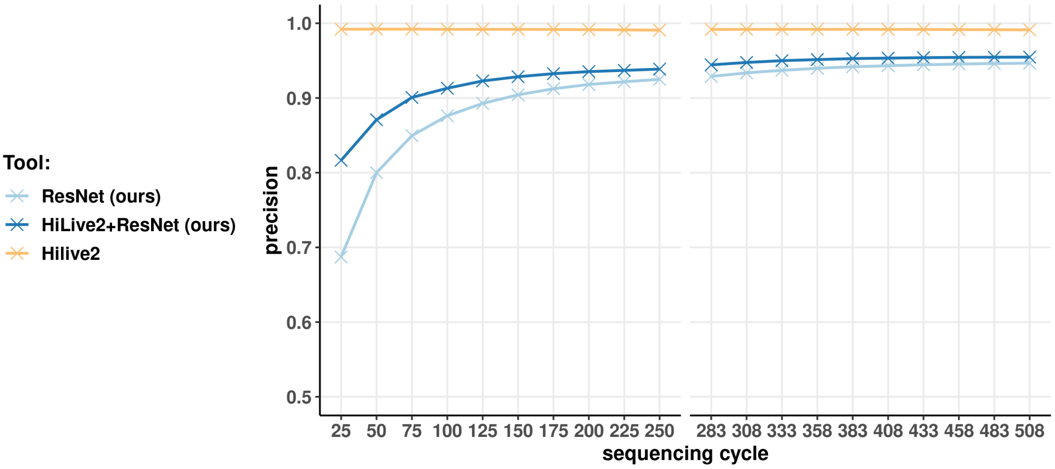


Figure S2: Precision for the viral dataset. We compared HiLive2's stable performance to the viral hybrid classifier and the ResNet alone, which achieved precision comparable to alignment-based approaches. The hybrid classifier crosses a 90% threshold at cycle 75 (90.1%), while never plunging below 80% even for the earliest cycles. What is more, all of the HiLive2-mapped reads are included in the hybrid classifier's predictions, so no information is lost by employing the extended approach. The high precision resulting from combining the real-time mapper with the deep learning classifier suggests that the associations of reads and a pathogenic phenotype are trustworthy even at the early stages of the sequencing run, getting even more reliable as more information is gained. Precision approaches HiLive2's, especially for the later cycles.

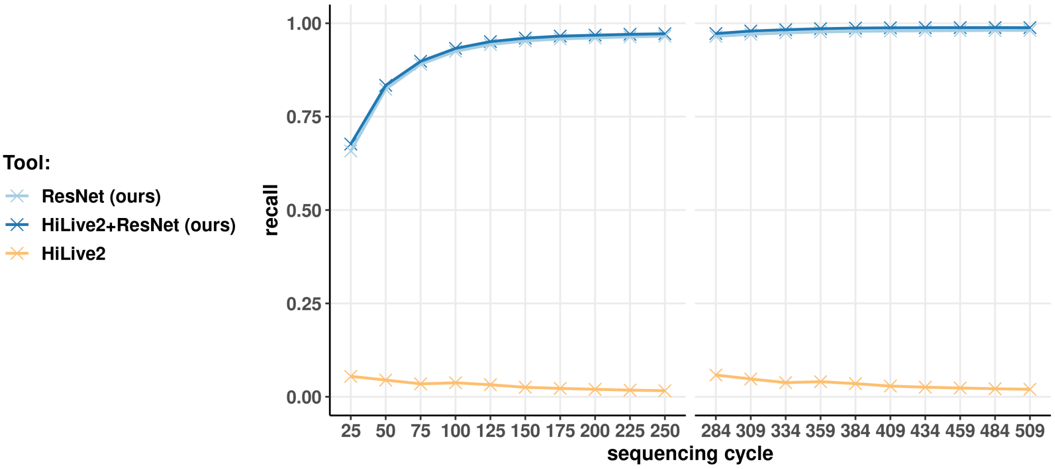


Figure S3: Recall (true positive rate) for the *S. aureus* sequencing run. As this is a pure pathogen sample, recall is equal to accuracy. A combination of HiLive2 and ResNet correctly identifies 98.8% of the read pairs after the last cycle, and 94.9% on average over the whole run. For this particular species, the performance of the ResNet itself is only marginally worse (98.1% after the last cycle and 94.0% on average).

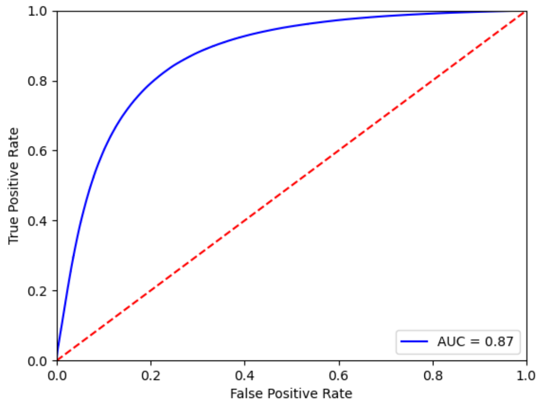


(a) ROC curve, bacteria

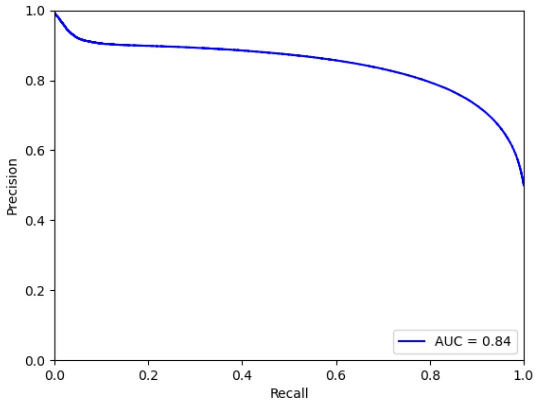


(b) PR curve, bacteria

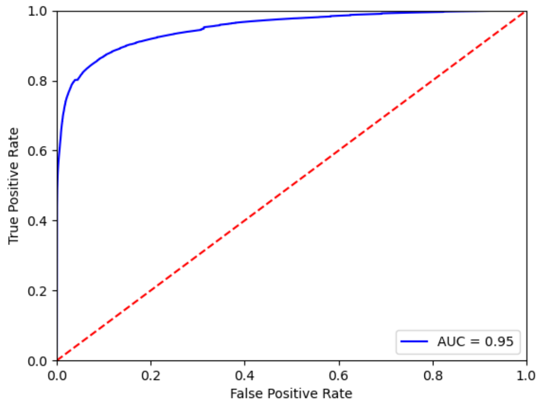


(c) ROC curve, viruses

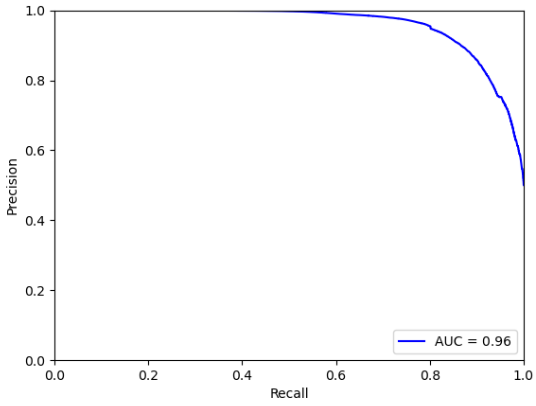


(d) PR curve, viruses

Figure S4: ROC and PR curves for Nanopore-trained ResNets evaluated on the bacterial and viral Nanopore test sets. Note that if a user wishes to retune the classification threshold according to custom optimality criteria, we would advise reevaluating the selected threshold on another, separate held-out dataset. Alternatively (if this not possible), one can select the threshold using the validation set (available at https://doi.org/10.5281/zenodo.4456857), and then perform the final evaluation with a fixed threshold on the test set.
